# VEZF1 facilitates pluripotency exit by regulating developmental transcriptional programs and CTCF occupancy

**DOI:** 10.64898/2026.08.13.744713

**Authors:** Isaiah K. Mensah, Ming He, Mudasir Zahoor, Sameer U. Khan, Martin L. Emerson, Hern J. Tan, Geordan D. Bolden, Sagar M. Utturkar, Humaira Gowher

## Abstract

Vascular Endothelial Zinc Finger 1 (VEZF1) is essential for embryonic development, but its role in pluripotent state transitions remains unclear. Previous work showed that *Vezf1^-/-^* ESCs exhibit impaired differentiation, reduced Dnmt3b expression, and genome-wide hypomethylation. Here, our systematic investigation shows that VEZF1 is required for pluripotency exit during ESC differentiation. *Vezf1^-/-^* ESCs fail to efficiently repress the pluripotency transcriptional program during differentiation, a defect that persists despite ectopic Dnmt3b expression. Genome-wide analysis revealed VEZF1 occupancy at regulatory regions of genes involved in several signaling pathways, including MAPK, as well as at some pluripotency-associated genes. Many VEZF1- bound MAPK genes showed reduced expression in undifferentiated *Vezf1^-/-^* ESCs, suggesting that VEZF1 activity contributes to the transcriptional competence required for efficient differentiation. VEZF1 loss also led to widespread acquisition of new CTCF sites associated with developmental signaling, a subset of which overlapped VEZF1-bound regulatory regions. CTCF depletion partially rescued the expression of several tested MAPK pathway genes, indicating that altered CTCF occupancy contributes to their reduced expression in *Vezf1^-/-^* ESCs. Together, our findings identify a novel function for VEZF1 in coordinating pluripotency exit with expression of pro-differentiation signaling genes and reveal a functional interplay between VEZF1 and CTCF during cell state transitions.

## Introduction

Pluripotency exit and differentiation are distinct but interconnected processes: the former terminates the pluripotent state, whereas the latter establishes lineage-specific identity (Li et al. 2020b). This transition requires coordinated repression of core pluripotency factors and activation of developmental signaling pathways, including MAPK signaling (Young 2011; Vega-Sendino et al. 2021). The mechanisms integrating transcriptional regulation and chromatin organization to establish competence for early developmental transition remain incompletely understood.

Murine embryonic stem cells (ESCs) can be maintained in medium supplemented with leukemia inhibitory factor (LIF), which activates JAK–STAT signaling to support pluripotency and self-renewal (Hirai et al. 2011). Under serum/LIF conditions, ESC cultures are heterogeneous and fluctuate between transcriptional states associated with naïve pluripotency and differentiation competence (Hayashi et al. 2008). The mitogen-activated protein kinase–extracellular signal-regulated kinase (MAPK–ERK) pathway plays a central role in destabilizing the naïve pluripotency network (Kunath et al. 2007; Silva and Smith 2008), and inhibition of MAPK–ERK signaling together with LIF efficiently maintains ESCs in the naïve state (Dunn et al. 2014). Conversely, activation of MAPK–ERK signaling promotes release from naïve pluripotency and progression toward differentiation. Downstream MAPK signaling promotes displacement of ERF from regulatory regions of naïve pluripotency genes, facilitating their repression, and contributes to induction of Dnmt3b and other differentiation-associated genes (Vega-Sendino et al. 2021). Despite the importance of MAPK signaling in this transition, considerably less is known about the transcriptional mechanisms that establish appropriate expression of MAPK pathway components in pluripotent cells.

VEZF1 (Vascular Endothelial Zinc Finger 1) is a zinc-finger transcription factor with established functions in embryonic and vascular development. Genetic ablation of Vezf1 in mice results in embryonic lethality associated with vascular defects (Kuhnert et al. 2005). VEZF1 contains multiple C2H2 zinc-finger domains that bind GC-rich DNA sequences, while recent studies suggest additional interactions with noncanonical nucleic acid structures, including G-quadruplexes and R-loops (Li et al. 2020a; Li et al. 2025). Our previous studies showed that VEZF1 binds to CpG-rich promoters genome-wide and influences transcription elongation through regulation of RNA polymerase II pausing (Gowher et al. 2012). We also found that reduced Dnmt3b expression in *Vezf1^-/-^* ESCs results in widespread genomic hypomethylation (Gowher et al. 2008). Additionally, we reported that *Vezf1^-/-^* ESCs retained alkaline phosphatase activity after differentiation (AlAbdi et al. 2018), suggesting that VEZF1’s role in pluripotency involves mechanisms other than regulation of DNMT3B and DNA methylation.

*Vezf1* was initially identified as an insulator-binding protein that co-localizes with CTCF at the chicken β-globin HS4 insulator, where it participates in both barrier and enhancer-blocking activities (Dickson et al. 2010; Huang et al. 2021). CTCF is a major architectural protein that organizes chromatin interactions, contributes to topologically associating domain (TAD) boundaries, and can constrain or facilitate enhancer–promoter communication (Ong and Corces 2014; Merkenschlager and Nora 2016). Dynamic changes in enhancer–promoter interactions and higher-order chromatin organization accompany changes in gene expression during ESC state transitions and lineage specification (Gaszner and Felsenfeld 2006; Lieberman-Aiden et al. 2009; Giles et al. 2010; Handoko et al. 2011; Kalhor et al. 2011; Dowen et al. 2013; Calderon et al. 2022; Ealo et al. 2024). Although both VEZF1 and CTCF have been implicated in transcriptional and chromatin regulation, whether VEZF1 influences CTCF occupancy in pluripotent cells and how this relationship relates to developmental gene regulation remain unknown.

In this study, we examined how VEZF1 controls the pluripotency and differentiation potential of ESCs. Using transcriptomic, genomic, and functional analyses, we examined how VEZF1 deficiency alters the transcriptional programs associated with developmental transitions. Gene expression analysis showed that *Vezf1*^-/-^ ESCs fail to appropriately repress pluripotency-associated genes post-differentiation, and this defect persists despite restoration of DNMT3B expression.

Genome-wide analyses reveal VEZF1 occupancy at both pluripotency-associated genes and genes within major developmental signaling pathways, including MAPK, WNT, and Hippo, with reduced expression of many VEZF1-bound MAPK genes in undifferentiated *Vezf1*^-/-^ ESCs. In parallel, loss of VEZF1 was accompanied by extensive de novo CTCF binding, including acquisition of CTCF at some VEZF1-bound regulatory regions. CTCF depletion in *Vezf1^-/-^* ESCs partially restored expression of the MAPK genes examined, indicating that increased CTCF occupancy contributes to their reduced expression following VEZF1 loss. Together, these findings identify a novel function for VEZF1 in coordinating repression of the pluripotency program with regulation of MAPK pathway genes and reveal a previously unrecognized functional relationship between VEZF1 and CTCF binding. The impaired induction of VEZF1 in embryonal carcinoma cells, together with its reduced expression in several human cancers, suggests that this regulatory function may also be relevant to persistent stem-like transcriptional states in cancer.

## Results

### VEZF1 is required for efficient repression of the pluripotency program during ESC differentiation

WT and *Vezf1^-/-^* ESCs were differentiated by removal of LIF to form embryoid bodies (EB’s) followed by addition of retinoic acid (RA) on D3 for 4 additional days as previously described (Mummery et al. 1990; Chatagnon et al. 2015) (**Supplemental Fig. S1A)**. Total RNA was isolated from undifferentiated (UD) and differentiated cells collected on days 3, 5, and 7, and the expression of pluripotency genes, *Oct4* and *Nanog*, was analyzed by RT-qPCR. These transcription factors are essential components of the pluripotency regulatory network in both mouse and human ESCs (Takahashi and Yamanaka 2006; Takahashi et al. 2007; Kellner and Kikyo 2010). As expected, differentiation of WT ESCs resulted in progressive downregulation of *Oct4* and *Nanog*. In contrast, *Vezf1^-/-^* cells exhibited incomplete repression of both genes throughout differentiation (**Fig. 1A)**. Persistent expression of OCT4 protein in differentiated *Vezf1^-/-^* cells was also confirmed by Western blotting **(Fig. 1B**). Notably, Vezf1 expression increased approximately 3–4-fold during differentiation **(Fig. 1C),** consistent with a role for VEZF1 during the transition from pluripotency toward differentiation.

**Figure 1.**
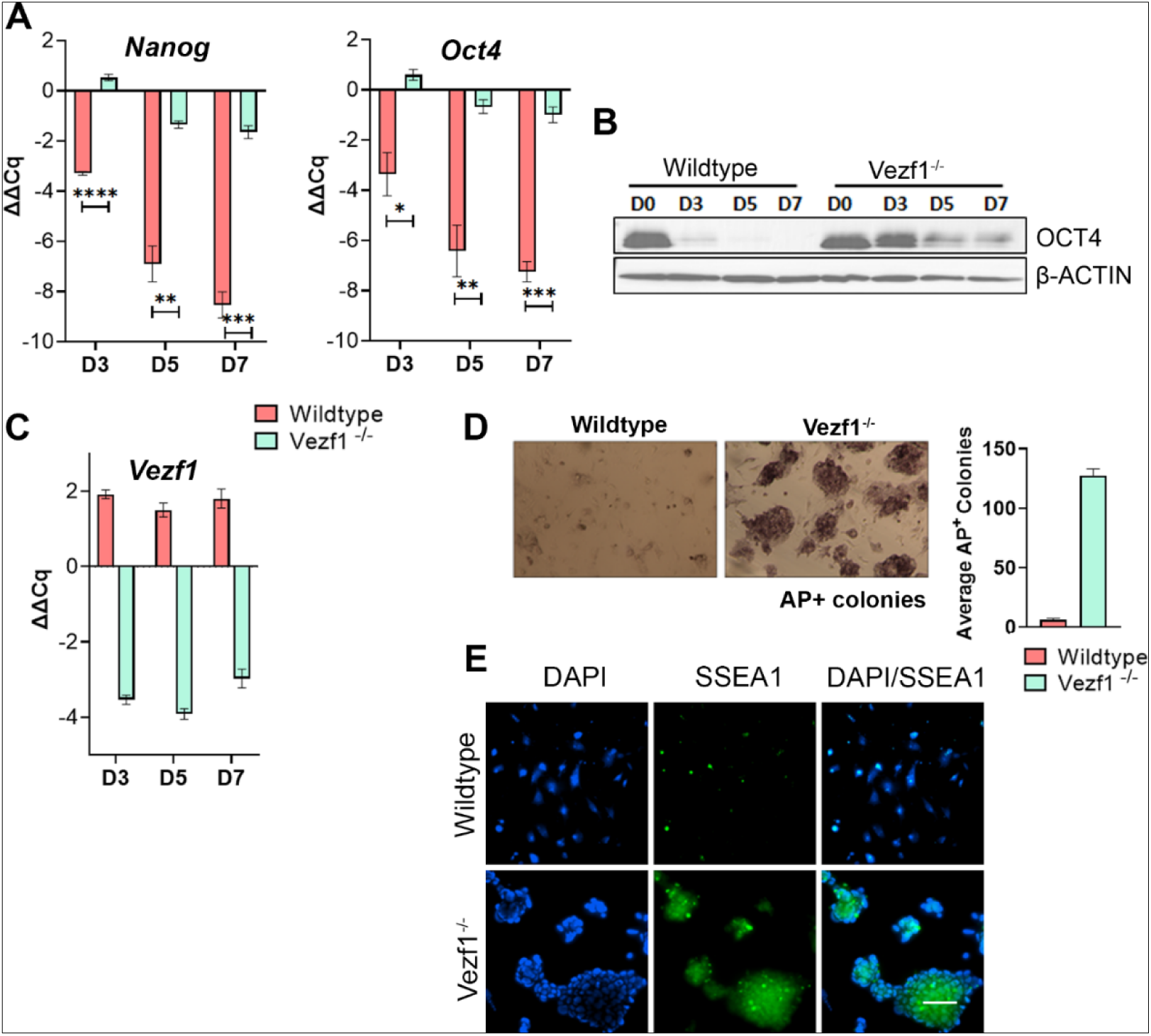
Retention of pluripotency in *Vezf1^-/-^* cells post-differentiation. (A, C) RT-qPCR showing the expression of (A) Oct4, Nanog, and (C) *Vezf1* in WT and *Vezf1*^-/-^ cells post-differentiation, relative to undifferentiated cells. Expression in undifferentiated cells is set to zero to illustrate both upregulation and downregulation. (B) Western blot showing the expression of OCT4 in WT and *Vezf1^-/-^* cells. β-ACTIN was used as a loading control. (D) Alkaline phosphatase stain of WT and *Vezf1*^-/-^ cells re-cultured with LIF following RA differentiation. The bar graph shows the number of colonies counted on a 100mm plate. (E) Immunofluorescence of SSEA1 in WT and *Vezf1*^-/-^ cells re-cultured with LIF after RA differentiation. The scale bar represents 100 µm. RA-retinoic acid; D3-D7 – days 3 to 7.

To determine whether persistent expression of pluripotency genes in *Vezf1^-/-^* cells was associated with an incomplete exit from the pluripotent state, WT and *Vezf1^-/-^* ESCs were differentiated in RA-containing medium for 5 days, then returned to LIF-containing ESC medium for an additional 2 days **(Supplemental Fig. S1B)**. Under these conditions, WT cells failed to survive and did not form alkaline phosphatase-positive colonies. In contrast, *Vezf1^-/-^* cells efficiently formed alkaline phosphatase-positive colonies **(Fig. 1D)** that also stained positive for the pluripotency-associated surface marker SSEA-1 **(Fig. 1E and Supplemental Fig. S1C)**, indicating that a population of *Vezf1^-/-^* cells retained pluripotent characteristics despite exposure to differentiation cues.

Together, these results demonstrate that VEZF1 is required for efficient downregulation of the pluripotency transcriptional program during ESC differentiation.

### DNMT3B restoration fails to rescue repression of the pluripotency transcriptional program

Previous studies have shown that *Vezf1^-/-^* ESCs express reduced levels of the DNA methyltransferase DNMT3B, compared to WT ESCs (Gowher et al. 2008). To determine whether reduced *Dnmt3b* expression underlies the failure of *Vezf1^-/-^*cells to repress the pluripotency program during differentiation, we generated stable *Vezf1^-/-^* cell lines expressing full-length Dnmt3b1 (*Vezf1^-/-+3b^*). Restoration of DNMT3B expression was confirmed at both the mRNA and protein levels (**Fig. 2A**). Ectopic DNMT3B expression rescued the genomic DNA hypomethylation previously observed in *Vezf1*^-/-^ ESCs, as assessed by methylation-sensitive restriction enzyme analysis (Gowher et al. 2008; Kinney et al. 2011) **(Supplemental Fig. S2A).** However, similar to *Vezf1^-/-^*cells, *Vezf1*^-/-+3b^ cells failed to efficiently repress *Oct4* and *Nanog* following differentiation (Fig. 2B), indicating that restoration of DNMT3B alone is insufficient to rescue the differentiation defect. (**Fig. 2B**).

**Figure 2.**
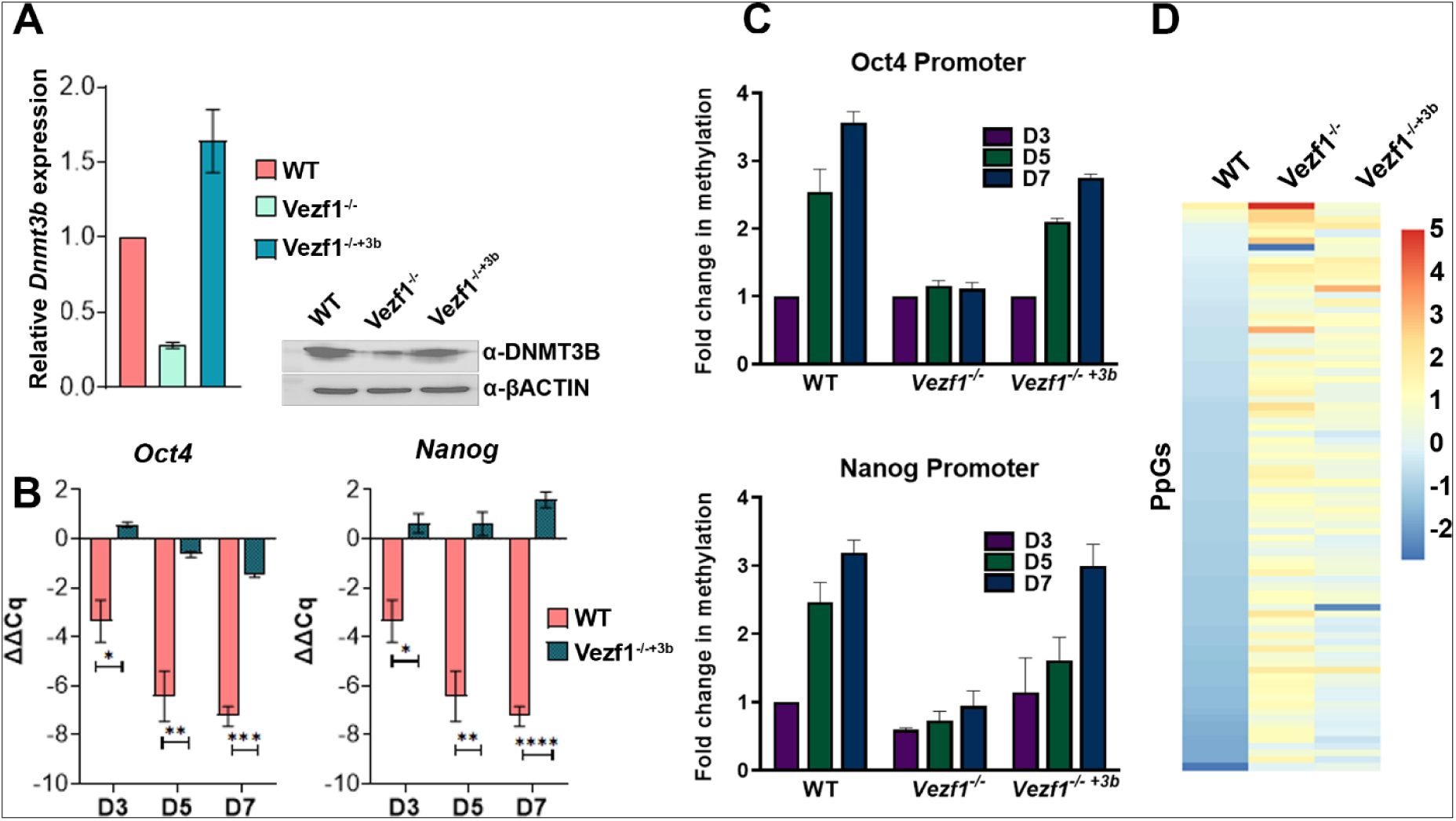
Ectopic DNMT3B expression fails to rescue pluripotency gene repression. (A) RT-qPCR showing Dnmt3b expression in WT, *Vezf1^-/-^* and *Vezf1^-/-+3b^* ESCs. Dnmt3b expression in *Vezf1^-/-^* mutants was normalized to wild-type cells, which was set to 1. Western blot showing the protein levels of DNMT3B in WT, *Vezf1^-/-^*, and *Vezf1^-/-+3b^* cells. β-actin was used as a loading control. (B) RT-qPCR showing the expression of Oct4 and Nanog in WT and *Vezf1^-/-^* + Dnmt3b cells. The change in Cq was normalized to undifferentiated cells, which were set to 0. (C) MD-qPCR showing the DNA methylation levels on cis-regulatory elements of pluripotency genes. (D) Heatmap showing pluripotency gene expression in wild-type versus *Vezf1^-/-^* mutants post-differentiation.

Because DNA methylation at pluripotency gene regulatory elements is cooperatively established by DNMT3A and DNMT3B (Hattori et al. 2004; Li et al. 2007; Kellner and Kikyo 2010; Shanak and Helms 2020), we next examined DNA methylation at *Oct4* and *Nanog* promoters in WT, *Vezf1^-/-^, and Vezf1^-/-+3b^*cells before and after differentiation using the methylation-dependent qPCR (MD-qPCR) method (Petell et al. 2017; Saha et al. 2020). As expected, promoter methylation was reduced in differentiated *Vezf1^-/-^* cells relative to WT cells. Re-expression of DNMT3B substantially restored DNA methylation at both promoters **(Fig. 2C)**. However, despite increased promoter methylation, *Oct4* and *Nanog* remained incompletely repressed, demonstrating that restoration of promoter DNA methylation alone is insufficient to silence the pluripotency program in the absence of VEZF1.

To determine whether this effect extended beyond Oct4 and Nanog, we performed RNA-Seq on undifferentiated and day 3-differentiated wild-type, *Vezf1^-/-^,* and *Vezf1^-/-+3b^* cells. Using a GFOLD cutoff of >1 and < -1, approximately 4,400 differentially expressed genes (DEGs) were identified during WT differentiation, whereas 9,000–11,000 DEGs were detected in *Vezf1^-/-^* and *Vezf1^-/-+3b^* cells **(Supplemental Table S1).** More than 50% of the DEGs were shared between *Vezf1^-/-^* and *Vezf1^-/-+3b^* cells, whereas only approximately half of the WT DEGs overlapped with either genotype (**Supplemental Fig. S2B**). Hierarchical clustering further showed that the transcriptional profiles of *Vezf1^-/-^* and *Vezf1^-/-+3b^* cells remained highly similar and distinct from WT cells (**Supplemental Fig. S2C**). Consistent with our RT-qPCR analyses, RNA-seq revealed robust repression of the pluripotency gene network during WT differentiation that was largely impaired in both *Vezf1^-/-^* and *Vezf1^-/-+3b^* cells. Although expression of a subset of pluripotency-associated genes was partially restored by DNMT3B re-expression, the global pluripotency transcriptional program remained largely unrepressed and closely resembled that of *Vezf1^-/-^* cells **(Fig. 2D)**.

Together, these findings demonstrate that VEZF1 regulates pluripotency exit through mechanisms distinct from its effects on DNMT3B expression and DNA methylation.

### Transient modulation of *Vezf1* expression alters the expression of pluripotency genes

To determine whether impaired repression of pluripotency-associated genes after VEZF1 loss reflects a requirement for VEZF1 rather than secondary adaptation to long-term culture, we generated transgenic ESCs carrying doxycycline (dox)-inducible *Vezf1* shRNA (*Vezf1^sh^*) (Supplemental Fig. S3A). Dox treatment consistently and significantly reduced *Vezf1* mRNA levels, accompanied by a corresponding decrease in VEZF1 protein levels, with no significant difference in uninduced (no dox) samples (Fig. 3A, B). We next differentiated *Vezf1^sh^* cells in the presence of doxycycline to maintain Vezf1 depletion throughout (Supplemental Fig. S3B) and measured *Oct4* and *Nanog* expression before and after differentiation. *Vezf1^sh^* cells also showed impaired downregulation of *Oct4* and *Nanog* compared to wild-type cells post-differentiation (Fig. 3C), although the effect was less pronounced than in constitutive *Vezf1^-/-^* cells (Fig. 2B, C). These findings demonstrate that acute VEZF1 depletion recapitulates the pluripotency gene-expression defect, arguing against prolonged culture or clonal adaptation as the basis of the phenotype.

**Figure 3.**
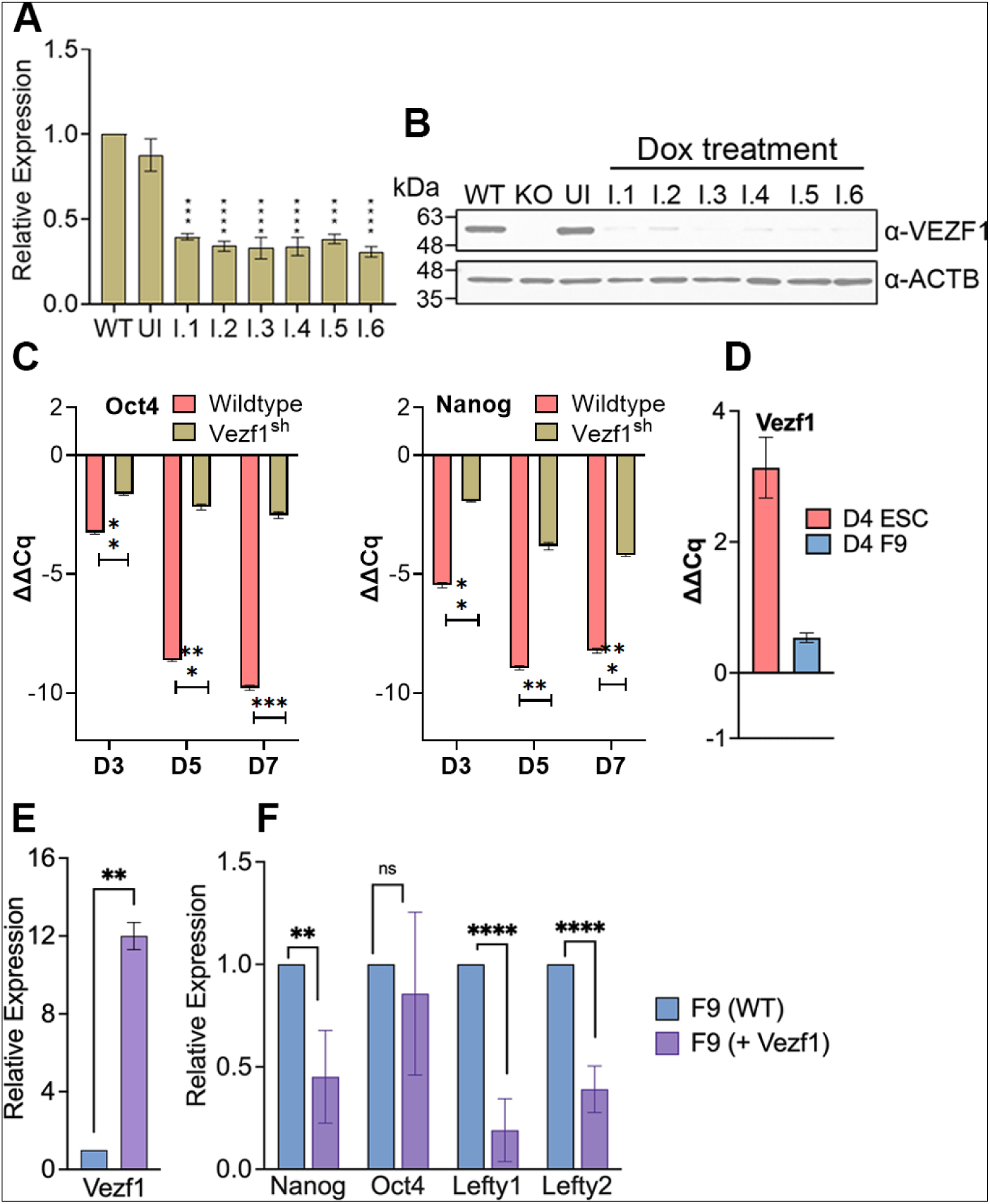
Modulation of VEZF1 expression alters pluripotency genes expression. (A) RT-qPCR of *Vezf1* in wild-type, uninduced (UI), and cells treated with doxycycline (I1-I6). (B) Western blot showing the expression of *Vezf1* in wild-type, uninduced, and *Vezf1^sh^* (knockdown) cells treated with doxycycline. Β-actin was used as a loading control. (C, D) RT-qPCR showing the expression of (C) Oct4 and Nanog in WT and *Vezf1^sh^* cells and (D) *Vezf1* in ESC and F9 embryonal carcinoma cells at D4 post-differentiation compared to undifferentiated cells. Cq values are normalized to Gapdh and compared to Cq values of undifferentiated cells, which are set to 0. (E, F) RT-qPCR showing fold change in expression of (E) *Vezf1* and (F) Pluripotency genes in F9 cells with ectopic *Vezf1* expression compared to WT F9 ECCs, where expression is set to 1.

To determine whether increased VEZF1 expression is sufficient to promote repression of pluripotency-associated genes, we next utilized F9 embryonal carcinoma cells (ECC), which fail to repress pluripotency genes following retinoic acid-induced differentiation **Supplemental Fig. S3C, S3D)** (Moore et al. 1985; Alonso et al. 1991; AlAbdi et al. 2020). Whereas *Vezf1* expression increased during ESC differentiation, its levels remained largely unchanged in differentiating F9 ECCs **(Fig. 3D, Supplemental Fig. S3E).** We therefore transiently overexpressed VEZF1 in F9 cells, which was confirmed by RT-qPCR (**Fig. 3E**). Ectopic VEZF1 expression significantly reduced the expression of pluripotency genes, *Nanog*, *Lefty1*, and *Lefty2*, whereas *Oct4* expression was not significantly altered (**Fig. 3F**). Collectively, these findings support a role for VEZF1 in repressing pluripotency-associated genes during early differentiation.

### VEZF1 occupies regulatory regions of pluripotency and developmental signaling genes

Given the established role of VEZF1 in transcriptional regulation (Gowher et al. 2008; Gowher et al. 2012), we performed ChIP-seq to identify its direct genomic targets in ESCs. Peak calling using MACS identified approximately 17,500 VEZF1 binding sites across the genome. Nearly 40% of VEZF1 peaks were localized to promoter regions, while approximately 32% were found in distal intergenic regions (**Supplemental Table S2**, **Fig. 4A, B**). Metagene analysis further demonstrated that VEZF1 binding was highly enriched around transcription start sites.

**Figure 4.**
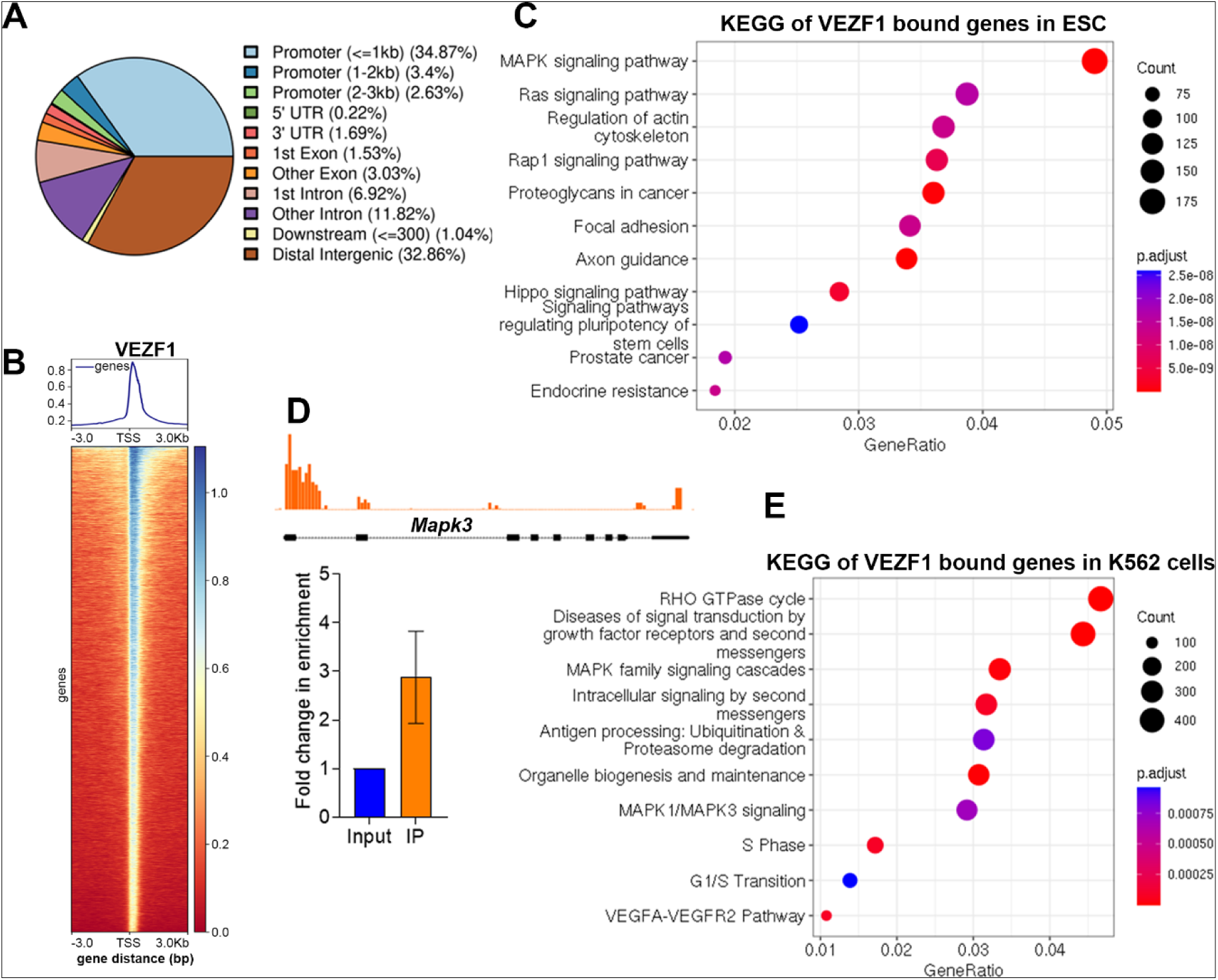
VEZF1 binds to genes in pro-differentiation signaling pathways. (A) Distribution of VEZF1 binding sites across the genome. (B) Profile plot and heatmap showing VEZF1 binding across the genome around the TSS. The scale bar shows the intensity of *Vezf1* enrichment. Profile plots and heat maps were generated using DeepTools. (C) KEGG pathway enrichment analysis of genes bound by VEZF1 in undifferentiated ESCs. (D) Example of VEZF1 ChIP-Seq data showing binding of VEZF1 to the MAPK3 promoter, and VEZF1 ChIP-qPCR validates the binding of VEZF1 to the MAPK3 promoter. (E) KEGG pathway enrichment analysis of genes bound by VEZF1 in K562, a human leukemia cell line.

To determine the biological processes directly associated with VEZF1 binding, we performed pathway enrichment analysis of VEZF1-bound genes. VEZF1 targets were significantly enriched for developmental signaling pathways, including MAPK, Ras, Rap1, Hippo, and focal adhesion pathways, as well as pathways regulating pluripotency of stem cells **(Fig. 4C)**. Thus, VEZF1 occupies genes associated with both the pluripotency network and signaling pathways that promote lineage commitment (**Supplemental Fig. S4A, Supplemental Table S2)**. VEZF1 occupancy at the Mapk3 promoter was independently validated by ChIP-qPCR (**Fig. 4D**).

To determine whether similar patterns of VEZF1 occupancy are observed in human cells, we analyzed publicly available VEZF1 ChIP-seq data from the human K562 cell line generated by the ENCODE Consortium. Similar to mouse ESCs, about 26% of Vezf1 binding sites (total 20,000) were localized to promoters and were enriched near transcription start sites, while approximately 23% occurred in distal intergenic regions (**Supplemental Fig. S4B, S4C**). Pathway enrichment analysis identified several signaling pathways among VEZF1-bound genes, including the RHO GTPase cycle, consistent with previous studies linking VEZF1 and RHO GTPases (Aitsebaomo et al. 2004; Gerald et al. 2013). Notably, MAPK signaling was also significantly enriched among VEZF1-bound genes, indicating that the association between VEZF1 occupancy and MAPK pathway genes is observed in both mouse ESCs and human K562 cells **(Fig. 4E**).

Finally, to characterize the chromatin environment associated with VEZF1 occupancy, we integrated the K562 VEZF1 ChIP-seq data with ENCODE datasets for RNA polymerase II Ser2 phosphorylation and histone modifications (Mzoughi et al. 2017). VEZF1 binding was strongly associated with regulatory regions marked by H3K27ac, H3K4me3, H3K4me1, and elongating RNA polymerase II, whereas little enrichment was observed at H3K27me3-marked regions **(Supplemental Fig. S4D**). These findings indicate that VEZF1 preferentially occupies transcriptionally active regulatory regions and identify developmental signaling and pluripotency-associated genes as prominent targets of VEZF1 occupancy.

### VEZF1 regulates the transcriptional state of MAPK signaling genes in ESCs

Having identified MAPK signaling as a major class of VEZF1-bound genes **(Fig. 4),** we next asked whether loss of VEZF1 altered the expression of these direct targets. We first extracted all VEZF1- bound genes from the RNA-Seq datasets of undifferentiated WT and *Vezf1^-/-^* ESCs. Differential expression analysis revealed a predominant reduction in transcript levels of VEZF1-bound genes in *Vezf1^-/-^* ESCs **(Fig. 5A)**, suggesting that VEZF1 contributes to maintaining their expression in undifferentiated ESCs.

**Figure 5.**
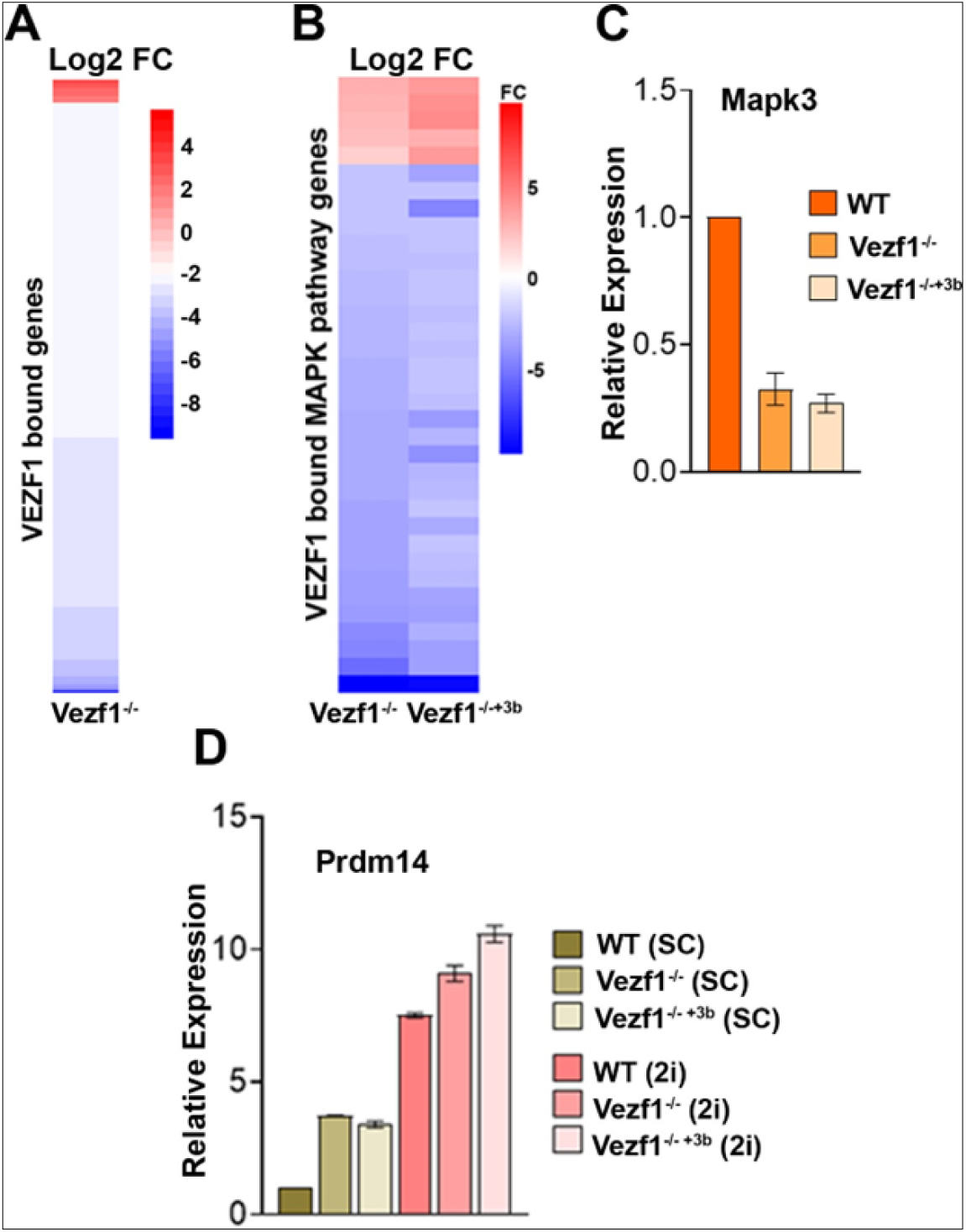
Vezf1-bound genes show reduced expression in *Vezf1^-/-^* compared to WT ESCs. (A, B) For undifferentiated ESCs, a heatmap showing the log2 fold change in expression in *Vezf1^- /-^* and *Vezf1^-/-+3b^*cells relative to WT (*WT/Vezf1^-/-^*) (*WT/Vezf1^-/-+3b^*) (A) for all differentially expressed genes and (B) for MAPK genes bound by VEZF1. (C) RT-qPCR of *Mapk3* gene expression in WT, *Vezf1^-/-^*, and *Vezf1^-/-+3b,^* showing reduced expression of *Mapk3* in the absence of *Vezf1*. (D) RT-qPCR for *Prdm14* expression in WT and *Vezf1^-/-^* mutants when cultured in serum-containing media (SCM) or 2i media (2i); FC = fold change.

We next focused on VEZF1-bound genes associated with the MAPK signaling pathway. Comparison of WT, *Vezf1^-/-^* and *Vezf1^-/-+3b^*showed that the majority of these genes were downregulated in both *Vezf1^-/-^* and *Vezf1^-/-+3b^* relative to WT cells **(Fig. 5B)**. Reduced expression of the representative target Mapk3 was independently confirmed by RT-qPCR **(Fig. 5C)**. Importantly, reduced expression did not reflect a general inability of *Vezf1^-/-^* cells to induce MAPK pathway genes post-differentiation. On Day 3, many MAPK-associated genes were induced, with a few showing greater induction in *Vezf1^-/-^* cells than in WT cells (**Supplemental Table S3)**. Consequently, some of the initially reduced VEZF1-bound MAPK genes approached or exceeded WT expression during differentiation, whereas others remained reduced. Thus, loss of VEZF1 does not uniformly attenuate the MAPK transcriptional response; instead, it alters the basal ESC-associated expression and subsequent transcriptional dynamics of multiple pathway components, potentially affecting their capacity to respond appropriately to differentiation cues. Reduced expression of VEZF1-bound MAPK genes is particularly notable because MAPK/ERK signaling is critical for the transition from naïve pluripotency toward differentiation (Mzoughi et al. 2017; Vega-Sendino et al. 2021). Consistent with this interpretation, serum-cultured *Vezf1^-/-^*ESCs exhibited significantly elevated expression of the naïve pluripotency marker Prdm14 compared with WT ESCs **(Fig. 5D)**, resembling the transcriptional state induced by pharmacological inhibition of MEK during 2i adaptation. As expected, culture under 2i conditions further increased Prdm14 expression in both WT and *Vezf1^-/-^* ESCs, and eliminated the difference between the two cell lines. A similar increase in Prdm14 expression was observed following acute VEZF1 depletion in doxycycline-inducible *Vezf1^sh^* ESCs **(Supplemental Fig. S5A**). Together, these findings indicate that VEZF1 regulates the transcriptional state of the MAPK signaling network, providing a transcriptional environment that supports efficient release from pluripotency.

To explore the potential relevance of VEZF1 beyond embryonic stem cells, we analyzed VEZF1 expression in human cancers using GEPIA2. VEZF1 expression was significantly reduced in adenoid cystic carcinoma (ACC), bladder urothelial carcinoma (BLCA), cervical squamous cell carcinoma (CESC), and uterine carcinosarcoma (UCS) **(Supplemental Fig. S5B)**. MAPK3 expression was also reduced in several of these tumor types **(Supplemental Fig. S5C),** and low VEZF1 expression was associated with poorer overall survival in UCS patients **(Supplemental Fig. S5D).** These observations suggest that reduced VEZF1 expression and disruption of associated developmental signaling networks may also occur in pathological contexts characterized by altered cellular differentiation.

### Loss of VEZF1 results in widespread acquisition of CTCF sites at developmental regulatory **regions**

To gain further insight into the mechanisms by which VEZF1 regulates gene expression, we performed HOMER motif analysis to identify DNA-binding factors whose consensus motifs are enriched in VEZF1-bound regions. The CTCF consensus motif was the most significantly enriched motif identified **(Supplemental Fig. S6A)**. This finding was particularly intriguing given previous reports describing co-occupancy of VEZF1 and CTCF at the HS4 insulator element of the chicken β-globin locus and at additional genomic regions in ESCs (Dickson et al. 2010). To investigate the relationship between VEZF1 and CTCF binding, we performed CTCF ChIP-Seq in WT and *Vezf1^- /-^* ESCs. Loss of VEZF1 resulted in approximately 22,000 additional CTCF binding sites accompanied by a modest increase in total CTCF protein levels in *Vezf1^-/-^* ESCs (**Supplemental Table S4 and Supplemental Fig. S6B, S6C**). Comparison of CTCF binding profiles revealed 28,933 shared sites, representing ∼ 85% of CTCF sites in WT cells and ∼ 51% of CTCF sites in *Vezf1^-/-^*cells (**Fig. 6A**). These shared sites remained highly stable across multiple overlap thresholds (**Supplemental Fig. S6D, S6E**). Genome-wide annotation further showed that most CTCF sites localized to distal intergenic regions (∼43% of total sites), whereas approximately 10% occurred within promoter regions (**Supplemental Fig. S6F, S6G**).

**Figure 6.**
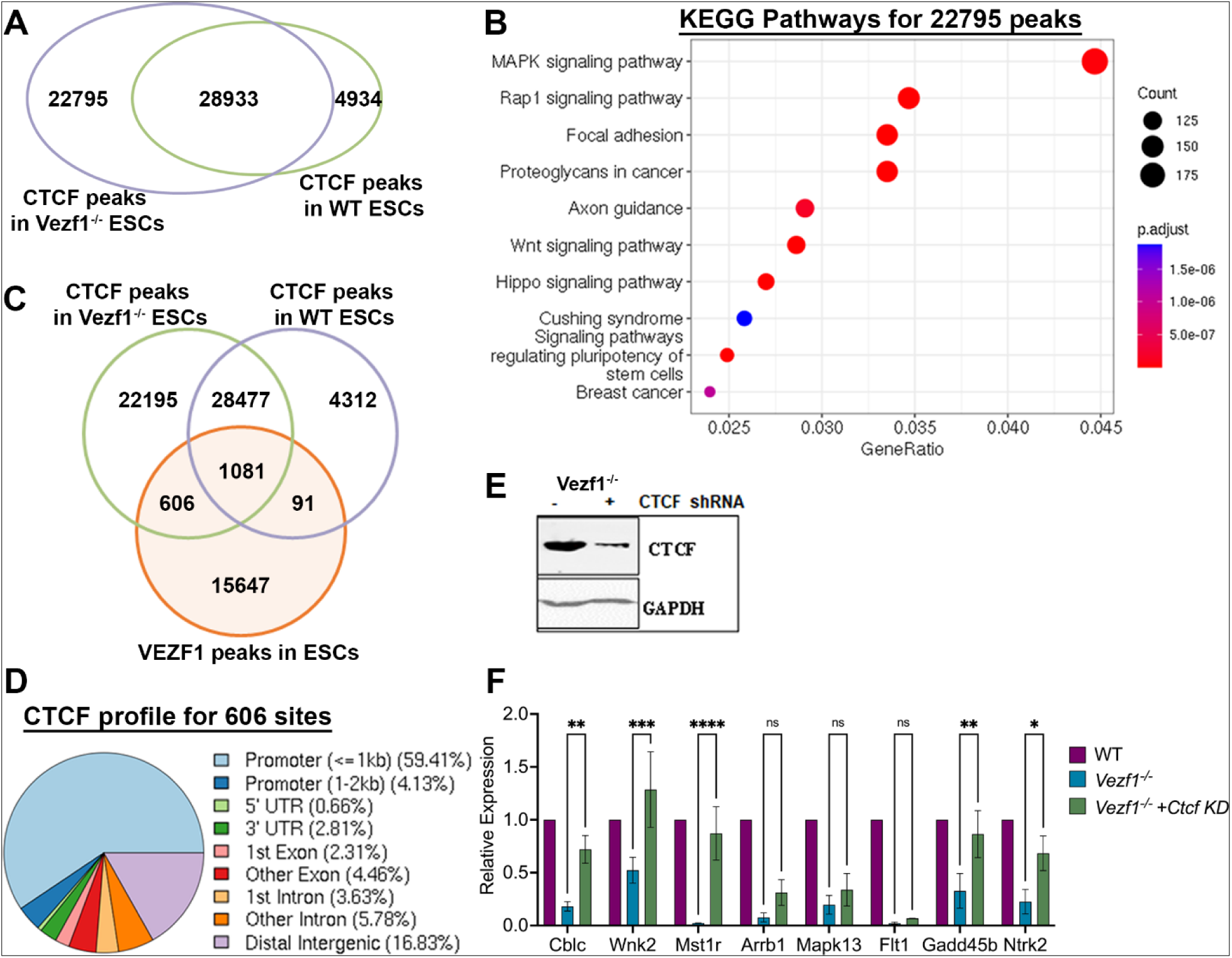
Increased CTCF occupancy in *Vezf1^-/-^* ESCs. (A) The Venn diagram shows the overlap of CTCF sites in WT and *Vezf1^-/-^* ESCs. (B) KEGG pathway enrichment analysis of CTCF sites that are unique in *Vezf1^-/-^* cells. (C) Venn diagram showing the overlap of VEZF1 sites and CTCF sites in WT and *Vezf1^-/-^* cells. (D) Genomic site distribution of CTCF sites in *Vezf1^-/-^* cells that overlap with VEZF1 sites. (E) Western Blot analysis showed reduced expression of CTCF in shRNA-transfected *Vezf1^-/-^* ESCs for 72 hrs. (F) RT-qPCR showing the expression of some MAPK pathway genes in *Vezf1^-/-^* ESCs with and without CTCF depletion. Error bars are SEM of triplicate data from two independent CTCF KD experiments.

Pathway enrichment analysis of CTCF sites uniquely acquired in *Vezf1^-/-^* cells identified significant enrichment for genes involved in MAPK, Hippo, and WNT, as well as the “Signaling pathways regulating pluripotency of stem cells” (**Fig. 6B**). Intersection of VEZF1 peaks with CTCF peaks identified 606 newly acquired CTCF sites within VEZF1-bound regions, approximately 60% of which were promoter-associated **(Fig. 6C, D)**. Genes associated with these sites were also enriched for MAPK signaling components **(Supplemental Fig. S6H)**.

To determine whether the increased CTCF occupancy observed in *Vezf1^-/-^*cells contributes to altered expression of MAPK-associated genes, we transiently depleted CTCF using shRNA. Efficient depletion of CTCF was confirmed by Western blot analysis (**Fig. 6E).** RT-qPCR analysis of selected MAPK genes, including both VEZF1-bound (*Arrb1, Mapk13, Flt1, Gadd45b*, and *Ntrk2)* and unbound targets (*Cblc, Wnk2*, and *Mst1r)*, showed that CTCF depletion shifted their expression toward wild-type levels, although the magnitude of recovery varied among genes (**Fig. 6F)**. These findings indicate that altered CTCF occupancy contributes to transcriptional changes following VEZF1 loss at both directly and indirectly regulated genes, while the incomplete recovery of VEZF1-bound genes is consistent with an additional direct contribution of VEZF1 to their regulation.

Together, these data show that VEZF1 loss is accompanied by widespread acquisition of new CTCF binding sites enriched near genes involved in pluripotency and pro-differentiation signaling pathways, a subset of which overlaps VEZF1-bound regulatory regions. Partial recovery of the MAPK genes examined following CTCF depletion establishes a functional consequence of altered CTCF occupancy and suggests that CTCF contributes to, but does not fully account for, the transcriptional effects of VEZF1 loss.

## Discussion

VEZF1 is a C2H2 zinc-finger transcription factor implicated in diverse developmental and physiological processes, including endothelial differentiation, angiogenesis, cardiovascular development, vascular integrity, and DNA methylation regulation (Katz et al. 2002; Kuhnert et al. 2005; Matsumoto et al. 2006; Gowher et al. 2008). However, the molecular mechanisms through which VEZF1 coordinates these processes remain poorly understood. Our previous work showed that *Vezf1^-/-^* ESCs tested positive for alkaline phosphatase after differentiating into endothelial cells, suggesting impaired transition from the pluripotent state (AlAbdi et al. 2018). Here, we identify VEZF1 as a key regulator of pluripotency exit in mouse embryonic stem cells. We demonstrate that VEZF1 regulates components of both the pluripotency transcriptional network and pro-differentiation signaling pathways, thereby facilitating the transition from the pluripotent to the differentiated state.

Loss of VEZF1 resulted in persistent expression of the pluripotency program and retention of pluripotent characteristics following differentiation. Genome-wide transcriptomic analyses further revealed that this phenotype extends beyond a few master regulators, with general retention of pluripotency-associated gene expression during differentiation of *Vezf1^-/-^* ESCs. Importantly, restoring DNMT3B expression in *Vezf1^-/-^* ESCs failed to rescue these defects, implicating a mechanism distinct from VEZF1-mediated regulation of DNMT3B and DNA methylation. Complementary studies using acute VEZF1 depletion and ectopic expression further supported a role for VEZF1 in repression of pluripotency-associated genes. Persistent activation of pluripotency-associated transcriptional programs is frequently associated with cancer stem-cell states (Astigiano et al. 2005; Yu et al. 2012; Zhao 2016; Shukla et al. 2017; Ou et al. 2018; Mahalaxmi et al. 2019). In several human tumor types, VEZF1 expression was reduced, with lower levels associated with poorer overall survival in uterine carcinosarcoma **(Supplemental Fig. S5C, S5E).** Together with our observations in F9 embryonal carcinoma cells, these findings suggest that reduced VEZF1 expression may be associated with persistent stem-like transcriptional states in cancer.

ChIP-seq revealed that VEZF1 occupies regulatory regions of genes involved in developmental signaling pathways, including MAPK, WNT, and Hippo, as well as a subset of pluripotency-associated genes. During differentiation, impaired repression was observed across the pluripotency gene network, affecting both VEZF1-bound and unbound genes. VEZF1 may therefore regulate this network through direct effects at a subset of occupied genes and secondary transcriptional effects that extend to unbound genes. Notably, most MAPK genes, including VEZF1-bound and unbound genes, were expressed at lower levels in undifferentiated *Vezf1^-/-^* ESCs, suggesting that VEZF1 establishes the basal expression of developmental signaling genes in pluripotent ESCs, thereby priming cells for an efficient response to differentiation cues. Although many MAPK genes approach wild-type expression levels by day 3, their reduced basal expression may compromise the earliest transcriptional response to differentiation cues, thereby compromising efficient repression of the pluripotency program. Interestingly, several tumor types with reduced VEZF1 expression also showed reduced MAPK3 expression **(Supplemental Fig. S5D)**, paralleling our observations in ESCs.

In addition to its effects on transcription, loss of VEZF1 markedly altered CTCF occupancy, with thousands of new CTCF-binding sites enriched at genes involved in developmental signaling and pluripotency. CTCF is a well-studied insulator protein that predominantly occupies intergenic regions and regulates enhancer–promoter interactions (Ong and Corces 2014), consistent with the distribution observed in our study. Despite CTCF’s context-dependent roles in gene regulation, promoter-associated CTCF binding has been linked to transcriptional repression (Filippova et al. 1996; Arnold et al. 2000; Dehingia et al. 2022). Our data show that many of the newly occupied CTCF sites in *Vezf1^-/-^* ESCs that overlap with VEZF1 binding are located in promoter regions. CTCF depletion partially restored expression of a subset of MAPK genes, providing functional evidence that altered CTCF binding contributes to the transcriptional changes associated with VEZF1 loss. The incomplete recovery suggests that CTCF represents one component of a more complex regulatory mechanism. Dynamic remodeling of CTCF occupancy accompanies lineage specification and contributes to the reorganization of chromatin architecture during developmental progression (Bonev et al. 2017; Nora et al. 2017). A similar relationship has been reported for ADNP, whose loss in ESCs results in widespread gain of CTCF occupancy, with consequences for chromatin organization during neural differentiation (Wulfridge et al. 2026). These parallel observations suggest that regulation of CTCF binding by distinct chromatin and transcriptional regulators may help establish a chromatin environment permissive to appropriate cell-state transitions.

Our findings establish VEZF1 as a regulator of pluripotency exit by supporting expression of MAPK signaling genes that contribute to transcriptional competence for differentiation. Beyond its effects on gene expression, the widespread acquisition of new CTCF sites upon VEZF1 loss together with the partial recovery of MAPK gene expression following CTCF depletion identifies a previously unrecognized relationship between VEZF1 and CTCF occupancy. Together, these findings provide a framework for understanding how VEZF1 coordinates transcriptional regulation with changes in CTCF binding and may help explain its diverse functions in embryonic development. Future studies will determine whether this relationship influences higher-order chromatin organization, lineage specification, cellular plasticity and disease.

## Materials and Methods

### ESC differentiation, 2i adaptation, and pluripotency exit assays

Retinoic acid differentiation: WT and Vezf1 mutant ESCs were differentiated as embryoid bodies (EBs) in suspension culture in ES medium without LIF (ES–LIF). Retinoic acid (RA; 1 µM) was added to D3 EBs, and the differentiation medium was replaced every other day; samples were collected on days 3, 5, and 7.

### 2i adaptation

Serum-cultured ESCs were washed to remove residual FBS and replated on 0.1% gelatin-coated plates in 2i medium containing 3 µM CHIR99021 and 1 µM PD0325901. Cells were passaged at least seven times before gene expression analysis.

### Pluripotency exit assay

WT and mutant ESCs were differentiated as EBs in ES–LIF medium at 1 × 10^6^ cells/mL. On day 3, 1 µM RA was added for 48 h. EBs were then dissociated and replated on 0.1% gelatin-coated plates in ES + LIF medium for 48 h. Cells were analyzed by alkaline phosphatase staining and SSEA-1 immunofluorescence. For alkaline phosphatase staining, cells were fixed with 1% formaldehyde for 5 min, quenched with 150 mM glycine, washed with PBS, and stained using an alkaline phosphatase staining kit according to the manufacturer’s instructions (Sigma, AB0300).

### Microscopy and Immunofluorescence

Brightfield images were acquired using a Zeiss microscope at 2.5× and 10× magnification, and relative EB size was quantified using ImageJ. For immunofluorescence, cells grown on gelatin-coated coverslips were fixed with 4% PFA for 15 min and incubated overnight at 4°C with primary antibodies in 5% milk/PBST (PBS, 0.4% Triton X-100). Cells were washed and incubated with Alexa Fluor 488-conjugated anti-mouse secondary antibody for 30 min at room temperature, mounted with DAPI-containing antifade medium, and imaged using a Leica DM6B fluorescence microscope.

### RNA purification and RT-qPCR

RNA from ESCs and differentiated cells was isolated using TRIzol reagent (Invitrogen, 15596026) according to the manufacturer’s instructions. The RNA samples were treated with DNase (Roche, 04716728001) for 2 hours at 37 °C to deplete them of contaminating genomic DNA and then repurified using a Quick-RNA™ Miniprep Plus Kit (Zymo Research, R1057). Next, 1 μg of RNA samples was used for cDNA synthesis using the Tetro cDNA Synthesis Kit (Bio line, BIO-65043). qPCR was performed using RT-qPCR kits (Thermo Scientific, AB-4104A) and primers (see primer list) according to the manufacturer’s instructions. Cq values were measured using the CFX Connect Real-Time PCR Detection System (Bio-Rad, 1855201). Gene expression was measured as ΔCt, defined as Cq (Target gene) - Cq (GAPDH or Β-Actin). Changes in gene expression are reported as fold change relative to control undifferentiated cells, which were set to 1.

### Genomic DNA purification and DNA methylation analysis

Genomic DNA was isolated by proteinase K digestion followed by phenol-chloroform extraction and RNase treatment. For methylation-dependent qPCR (MD-qPCR), purified DNA was digested overnight with FspEI (NEB, R0662S), repurified, and quantified using PicoGreen (Life Technologies, P11495). Equal amounts of DNA were analyzed by qPCR, and methylation was quantified as relative fold change in Cq. Primers were described previously (AlAbdi et al. 2020). Experiments included three biological and three technical replicates.

For global DNA methylation analysis, genomic DNA from WT, *Vezf1^-/-^*and *Vezf1^-/-+3b^* ESCs was digested with the methylation-sensitive restriction enzyme HpaII or its methylation-insensitive isoschizomer, MspI. Digestion patterns were analyzed to compare genomic methylation among the three cell lines.

### Chromatin immunoprecipitation

Cells were cross-linked with 1% formaldehyde for 5 min and quenched with 150 mM glycine. Nuclei were isolated by sequential washes with Triton X-100, high-salt, and TEG buffers and resuspended in shearing buffer (TEG, 100 mM NaCl, 0.1% sodium deoxycholate, 0.5% IGEPAL). Chromatin was sonicated using a Bioruptor Plus (Diagenode, B01020001) and cleared by centrifugation. Protein A/G magnetic beads (Life Technologies, 10002D/10004D) were incubated with 8 µg antibody and subsequently with 8 µg sheared chromatin overnight. Immunoprecipitated chromatin was eluted, cross-links were reversed at 65°C, and samples were treated with RNase and Proteinase K. DNA was purified by phenol-chloroform extraction and ethanol precipitation, quantified using PicoGreen, and used for qPCR and sequencing.

### ChIP-seq Data Processing and Analysis

Raw ChIP-seq data were generated using the Illumina sequencing platform. Data quality control was performed using fastp (v0.23.2) and FastQC (v0.12.1). Adapter sequences and low-quality bases (Phred quality score < 30) were removed, and a minimum read length of 50 bp was enforced. Sequencing quality was assessed before and after trimming using FastQC. High-quality reads were aligned to the Mus musculus reference genome (GRCm38) using Bowtie2 (v2.5.1). Only reads with a mapping quality score ≥10 were retained for downstream analyses. Normalized coverage tracks were generated in BigWig format using deepTools (v3.5.5) bamCoverage with CPM normalization. Peak calling was performed using two independent approaches. Peak calling was performed using MACS2 (v2.2.7.1) in paired-end mode with a false discovery rate (FDR) threshold of 0.05. Peaks identified by both approaches were annotated to nearby genes and genomic features using the ChIPseeker (v1.38.0) R package. Differential binding analysis was conducted using the DiffBind (v3.12.0) R package. Overlap among peak sets was visualized as Venn diagrams using DiffBind. Genome-wide signal enrichment profiles and heatmaps were generated using deepTools.

### RNA-seq Data Processing and Analysis

Raw RNA-seq data generated on the Illumina platform were processed using fastp to remove adapter contamination and low-quality bases (Phred quality score < 30). Reads shorter than 50 bp after trimming were discarded. Filtered reads were aligned to the Mus musculus reference genome (GRCm38) using the STAR (v2.7.11b) aligner. Gene-level quantification was performed using featureCounts from the subread package (v2.0.1) followed by differential expression analysis using edgeR (v4.0.16). For RNA-seq experiments lacking biological replicates, differential gene expression analysis was performed using GFOLD (v1.1.4), which estimates reliable fold changes in the absence of replicates. Gene-level read counts and subsequent pairwise differential expression analysis between treatment and control conditions were performed using the GFOLD-count and GFOLD-diff utility, respectively. The resulting GFOLD values were used as a robust estimate of gene up- or down-regulation. Genes with GFOLD values ≥1 or ≤−1 were considered differentially expressed and were retained for downstream analyses.

### Functional Enrichment and Pathway Analysis

Functional enrichment analysis of differentially expressed genes and specific peak-set-associated genes was performed using the clusterProfiler (v4.10.1) R package against the Kyoto Encyclopedia of Genes and Genomes (KEGG) database.

### Data Visualization

Data visualization was performed using R (version 4.3.3). Heatmaps were generated using the pheatmap (v1.0.12) and ComplexHeatmap (v2.18.0) packages. Additional visualizations were generated using ggplot2 (v3.5.1) and ggpubr (v0.6.0) R packages. ChIP-seq signal profiles and enrichment heatmaps were generated using the deepTools suite. UCSC browser images were generated using the mm10 genome.

### Reference Genome and Annotation Resource

All sequencing analyses were performed using the Mus musculus (GRCm38) reference genome and the corresponding gene annotation in GTF format, downloaded from the Ensembl database.

### Statistical Analysis

Statistical analyses were performed using R version 4.3.3. For ChIP-seq analyses, statistical significance was determined using the default thresholds of the respective peak-calling tools, unless otherwise stated. For RNA-seq analyses, edgeR results were filtered using a false discovery rate (FDR) cutoff of 0.05, while experiments lacking biological replicates were analyzed using GFOLD, with genes having GFOLD values ≥1 or ≤−1 considered differentially expressed. Pathway enrichment analysis was performed with a significance cutoff of p-value < 0.05.

### Data Sources and Availability

Pluripotency-associated gene lists were derived from published datasets reported (Whyte et al. 2012), and additional stem cell regulatory datasets were obtained from (Mallon et al. 2013). Enhancer annotations were obtained from the dataset reported by (Whyte et al. 2012). Raw sequencing data will be deposited in the NCBI Gene Expression Omnibus (GEO) and Sequence Read Archive (SRA).

## Acknowledgments

We acknowledge the Purdue Institute for Cancer Research and the associated NCI grant (grant P30CA023168) as well as support from the Walther Cancer Foundation.

## Author Contributions

H.G. conceived and supervised the study, designed experiments, analyzed and interpreted data, and wrote the manuscript. I.K.M. and M.Z. performed experiments, analyzed data, and prepared figures. M.H., S.U.K., M.L.E, H.J.T., and G.D.B performed experiments, and S.M.U performed bioinformatic data analysis.

## References

Aitsebaomo J, Wennerberg K, Der CJ, Zhang C, Kedar V, Moser M, Kingsley-Kallesen ML, Zeng GQ, Patterson C. 2004. p68RacGAP is a novel GTPase-activating protein that interacts with vascular endothelial zinc finger-1 and modulates endothelial cell capillary formation. J Biol Chem 279: 17963–17972.

AlAbdi L, He M, Yang Q, Norvil AB, Gowher H. 2018. The transcription factor Vezf1 represses the expression of the antiangiogenic factor Cited2 in endothelial cells. The Journal of biological chemistry 293: 11109–11118.

AlAbdi L, Saha D, He M, Dar MS, Utturkar SM, Sudyanti PA, McCune S, Spears BH, Breedlove JA, Lanman NA et al. 2020. Oct4-Mediated Inhibition of Lsd1 Activity Promotes the Active and Primed State of Pluripotency Enhancers. Cell reports 30: 1478–1490 e1476.

Alonso A, Breuer B, Steuer B, Fischer J. 1991. The F9-EC cell line as a model for the analysis of differentiation. Int J Dev Biol 35: 389–397.

Arnold R, Mäueler W, Bassili G, Lutz M, Burke L, Epplen TJ, Renkawitz R. 2000. The insulator protein CTCF represses transcription on binding to the (gt)(22)(ga)(15) microsatellite in intron 2 of the HLA- DRB1(*)0401 gene. Gene 253: 209–214.

Astigiano S, Damonte P, Fossati S, Boni L, Barbieri O. 2005. Fate of embryonal carcinoma cells injected into postimplantation mouse embryos. Differentiation 73: 484–490.

Bonev B, Mendelson Cohen N, Szabo Q, Fritsch L, Papadopoulos GL, Lubling Y, Xu X, Lv X, Hugnot JP, Tanay A et al. 2017. Multiscale 3D Genome Rewiring during Mouse Neural Development. Cell 171: 557–572.e524.

Calderon L, Weiss FD, Beagan JA, Oliveira MS, Georgieva R, Wang YF, Carroll TS, Dharmalingam G, Gong W, Tossell K et al. 2022. Cohesin-dependence of neuronal gene expression relates to chromatin loop length. Elife 11.

Chatagnon A, Veber P, Morin V, Bedo J, Triqueneaux G, Sémon M, Laudet V, d’Alché-Buc F, Benoit G. 2015. RAR/RXR binding dynamics distinguish pluripotency from differentiation associated cis-regulatory elements. Nucleic acids research 43: 4833–4854.

Dehingia B, Milewska M, Janowski M, Pękowska A. 2022. CTCF shapes chromatin structure and gene expression in health and disease. EMBO reports 23: e55146.

Dickson J, Gowher H, Strogantsev R, Gaszner M, Hair A, Felsenfeld G, West AG. 2010. VEZF1 elements mediate protection from DNA methylation. PLoS genetics 6: e1000804.

Dowen JM, Bilodeau S, Orlando DA, Hubner MR, Abraham BJ, Spector DL, Young RA. 2013. Multiple structural maintenance of chromosome complexes at transcriptional regulatory elements. Stem Cell Reports 1: 371–378.

Dunn SJ, Martello G, Yordanov B, Emmott S, Smith AG. 2014. Defining an essential transcription factor program for naïve pluripotency. Science 344: 1156–1160.

Ealo T, Sanchez-Gaya V, Respuela P, Muñoz-San Martín M, Martin-Batista E, Haro E, Rada-Iglesias A. 2024. Cooperative insulation of regulatory domains by CTCF-dependent physical insulation and promoter competition. Nature communications 15: 7258.

Filippova GN, Fagerlie S, Klenova EM, Myers C, Dehner Y, Goodwin G, Neiman PE, Collins SJ, Lobanenkov VV. 1996. An exceptionally conserved transcriptional repressor, CTCF, employs different combinations of zinc fingers to bind diverged promoter sequences of avian and mammalian c-myc oncogenes. Molecular and cellular biology 16: 2802–2813.

Gaszner M, Felsenfeld G. 2006. Insulators: exploiting transcriptional and epigenetic mechanisms. Nat Rev Genet 7: 703–713.

Gerald D, Adini I, Shechter S, Perruzzi C, Varnau J, Hopkins B, Kazerounian S, Kurschat P, Blachon S, Khedkar S et al. 2013. RhoB controls coordination of adult angiogenesis and lymphangiogenesis following injury by regulating VEZF1-mediated transcription. Nature communications 4: 2824.

Giles KE, Gowher H, Ghirlando R, Jin C, Felsenfeld G. 2010. Chromatin boundaries, insulators, and long-range interactions in the nucleus. Cold Spring Harbor symposia on quantitative biology 75: 79–85.

Gowher H, Brick K, Camerini-Otero RD, Felsenfeld G. 2012. Vezf1 protein binding sites genome-wide are associated with pausing of elongating RNA polymerase II. Proceedings of the National Academy of Sciences of the United States of America 109: 2370–2375.

Gowher H, Stuhlmann H, Felsenfeld G. 2008. Vezf1 regulates genomic DNA methylation through its effects on expression of DNA methyltransferase Dnmt3b. Genes & development 22: 2075–2084.

Handoko L, Xu H, Li G, Ngan CY, Chew E, Schnapp M, Lee CW, Ye C, Ping JL, Mulawadi F et al. 2011. CTCF-mediated functional chromatin interactome in pluripotent cells. Nat Genet 43: 630–638.

Hattori N, Nishino K, Ko YG, Ohgane J, Tanaka S, Shiota K. 2004. Epigenetic control of mouse Oct-4 gene expression in embryonic stem cells and trophoblast stem cells. J Biol Chem 279: 17063–17069.

Hayashi K, de Sousa Lopes SMC, Tang F, Lao K, Surani MA. 2008. Dynamic equilibrium and heterogeneity of mouse pluripotent stem cells with distinct functional and epigenetic states. Cell Stem Cell 3: 391–401.

Hirai H, Karian P, Kikyo N. 2011. Regulation of embryonic stem cell self-renewal and pluripotency by leukaemia inhibitory factor. Biochem J 438: 11–23.

Huang H, Zhu Q, Jussila A, Han Y, Bintu B, Kern C, Conte M, Zhang Y, Bianco S, Chiariello AM et al. 2021. CTCF mediates dosage- and sequence-context-dependent transcriptional insulation by forming local chromatin domains. Nat Genet 53: 1064–1074.

Kalhor R, Tjong H, Jayathilaka N, Alber F, Chen L. 2011. Genome architectures revealed by tethered chromosome conformation capture and population-based modeling. Nat Biotechnol 30: 90–98.

Katz JP, Perreault N, Goldstein BG, Lee CS, Labosky PA, Yang VW, Kaestner KH. 2002. The zinc-finger transcription factor Klf4 is required for terminal differentiation of goblet cells in the colon. Development 129: 2619–2628.

Kellner S, Kikyo N. 2010. Transcriptional regulation of the Oct4 gene, a master gene for pluripotency. Histol Histopathol 25: 405–412.

Kinney SM, Chin HG, Vaisvila R, Bitinaite J, Zheng Y, Estève PO, Feng S, Stroud H, Jacobsen SE, Pradhan S. 2011. Tissue-specific distribution and dynamic changes of 5-hydroxymethylcytosine in mammalian genomes. J Biol Chem 286: 24685–24693.

Kuhnert F, Campagnolo L, Xiong JW, Lemons D, Fitch MJ, Zou Z, Kiosses WB, Gardner H, Stuhlmann H. 2005. Dosage-dependent requirement for mouse Vezf1 in vascular system development. Dev Biol 283: 140–156.

Kunath T, Saba-El-Leil MK, Almousailleakh M, Wray J, Meloche S, Smith A. 2007. FGF stimulation of the Erk1/2 signalling cascade triggers transition of pluripotent embryonic stem cells from self-renewal to lineage commitment. Development 134: 2895–2902.

Li JY, Pu MT, Hirasawa R, Li BZ, Huang YN, Zeng R, Jing NH, Chen T, Li E, Sasaki H et al. 2007. Synergistic function of DNA methyltransferases Dnmt3a and Dnmt3b in the methylation of Oct4 and Nanog. Molecular and cellular biology 27: 8748–8759.

Li L, Williams P, Gao Z, Wang Y. 2020a. VEZF1-guanine quadruplex DNA interaction regulates alternative polyadenylation and detyrosinase activity of VASH1. Nucleic acids research 48: 11994–12003.

Li QV, Rosen BP, Huangfu D. 2020b. Decoding pluripotency: Genetic screens to interrogate the acquisition, maintenance, and exit of pluripotency. Wiley Interdiscip Rev Syst Biol Med 12: e1464.

Li Y, Sheng Y, Di C, Yao H. 2025. Base-pair resolution reveals clustered R-loops and DNA damage-susceptible R-loops. Mol Cell 85: 1686–1702 e1685.

Lieberman-Aiden E, van Berkum NL, Williams L, Imakaev M, Ragoczy T, Telling A, Amit I, Lajoie BR, Sabo PJ, Dorschner MO et al. 2009. Comprehensive mapping of long-range interactions reveals folding principles of the human genome. Science 326: 289–293.

Mahalaxmi I, Devi SM, Kaavya J, Arul N, Balachandar V, Santhy KS. 2019. New insight into NANOG: A novel therapeutic target for ovarian cancer (OC). Eur J Pharmacol 852: 51–57.

Mallon BS, Chenoweth JG, Johnson KR, Hamilton RS, Tesar PJ, Yavatkar AS, Tyson LJ, Park K, Chen KG, Fann YC et al. 2013. StemCellDB: the human pluripotent stem cell database at the National Institutes of Health. Stem Cell Res 10: 57–66.

Matsumoto N, Kubo A, Liu H, Akita K, Laub F, Ramirez F, Keller G, Friedman SL. 2006. Developmental regulation of yolk sac hematopoiesis by Kruppel-like factor 6. Blood 107: 1357–1365.

Merkenschlager M, Nora EP. 2016. CTCF and Cohesin in Genome Folding and Transcriptional Gene Regulation. Annu Rev Genomics Hum Genet 17: 17–43.

Moore EE, Moritz EA, Mitra NS. 1985. A variant F9 embryonal carcinoma cell line which undergoes incomplete differentiation in retinoic acid. Cancer Res 45: 4387–4396.

Mummery CL, Feyen A, Freund E, Shen S. 1990. Characteristics of embryonic stem cell differentiation: a comparison with two embryonal carcinoma cell lines. Cell Differ Dev 30: 195–206.

Mzoughi S, Zhang J, Hequet D, Teo SX, Fang H, Xing QR, Bezzi M, Seah MKY, Ong SLM, Shin EM et al. 2017. PRDM15 safeguards naive pluripotency by transcriptionally regulating WNT and MAPK-ERK signaling. Nat Genet 49: 1354–1363.

Nora EP, Goloborodko A, Valton AL, Gibcus JH, Uebersohn A, Abdennur N, Dekker J, Mirny LA, Bruneau BG. 2017. Targeted Degradation of CTCF Decouples Local Insulation of Chromosome Domains from Genomic Compartmentalization. Cell 169: 930–944.e922.

Ong CT, Corces VG. 2014. CTCF: an architectural protein bridging genome topology and function. Nat Rev Genet 15: 234–246.

Ou M, Li S, Tang L. 2018. PRDM14: A Potential Target for Cancer Therapy. Curr Cancer Drug Targets 18: 945–956.

Petell CJ, Loiseau G, Gandy R, Pradhan S, Gowher H. 2017. A refined DNA methylation detection method using MspJI coupled quantitative PCR. Analytical biochemistry 533: 1–9.

Saha D, Norvil AB, Lanman NA, Gowher H. 2020. Simplified MethylRAD Sequencing to Detect Changes in DNA Methylation at Enhancer Elements in Differentiating Embryonic Stem Cells. Epigenomes 4.

Shanak S, Helms V. 2020. DNA methylation and the core pluripotency network. Dev Biol 464: 145–160.

Shukla V, Rao M, Zhang H, Beers J, Wangsa D, Buishand FO, Wang Y, Yu Z, Stevenson HS, Reardon ES et al. 2017. ASXL3 Is a Novel Pluripotency Factor in Human Respiratory Epithelial Cells and a Potential Therapeutic Target in Small Cell Lung Cancer. Cancer Res 77: 6267–6281.

Silva J, Smith A. 2008. Capturing pluripotency. Cell 132: 532–536.

Takahashi K, Tanabe K, Ohnuki M, Narita M, Ichisaka T, Tomoda K, Yamanaka S. 2007. Induction of pluripotent stem cells from adult human fibroblasts by defined factors. Cell 131: 861–872.

Takahashi K, Yamanaka S. 2006. Induction of pluripotent stem cells from mouse embryonic and adult fibroblast cultures by defined factors. Cell 126: 663–676.

Vega-Sendino M, Olbrich T, Tillo D, Tran AD, Domingo CN, Franco M, FitzGerald PC, Kruhlak MJ, Ruiz S. 2021. The ETS transcription factor ERF controls the exit from the naïve pluripotent state in a MAPK-dependent manner. Sci Adv 7: eabg8306.

Whyte WA, Bilodeau S, Orlando DA, Hoke HA, Frampton GM, Foster CT, Cowley SM, Young RA. 2012. Enhancer decommissioning by LSD1 during embryonic stem cell differentiation. Nature 482: 221–225.

Wulfridge P, Rell N, Doherty J, Fang K-C, Lynskey ML, Sarma K. 2026. ADNP regulates chromatin architecture and lineage fidelity during neural differentiation. PLOS Genetics 22: e1012081.

Young RA. 2011. Control of the embryonic stem cell state. Cell 144: 940–954.

Yu Z, Pestell TG, Lisanti MP, Pestell RG. 2012. Cancer stem cells. Int J Biochem Cell Biol 44: 2144–2151.

Zhao J. 2016. Cancer stem cells and chemoresistance: The smartest survives the raid. Pharmacol Ther 160: 145–158.

