## Supplemental Figures for "VEZF1 facilitates pluripotency exit by regulating developmental transcriptional programs and CTCF occupancy"

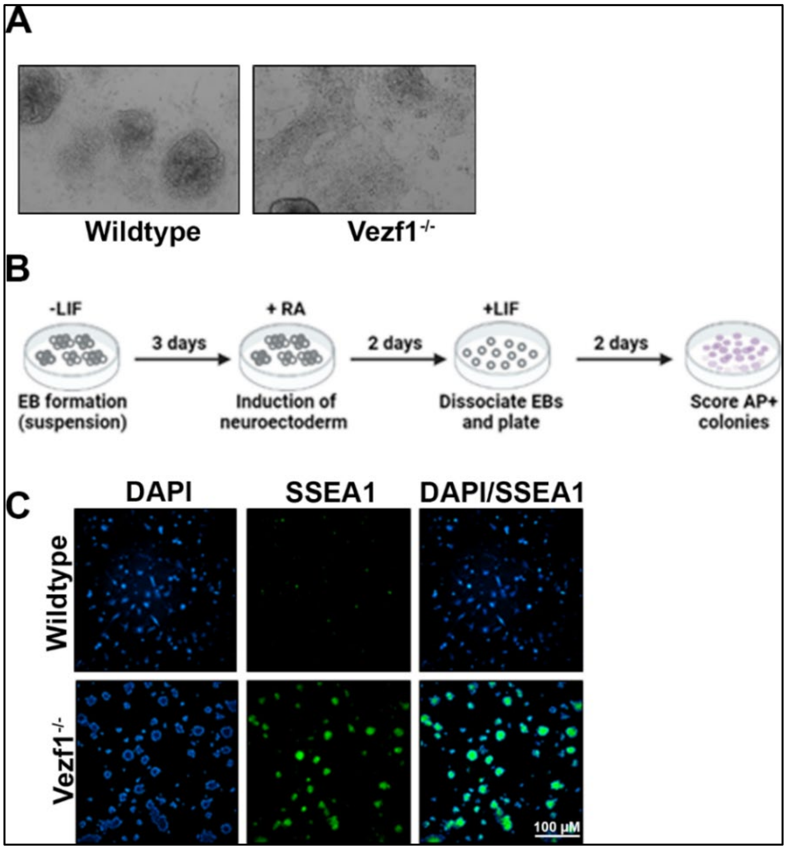


**Figure S1.** ***Vezf1***$^{-/-}$ **cells do not exit pluripotency post-differentiation**.

(A) Brightfield images of WT and *Vezf1*$^{-/-}$ cells at day 7 post-differentiation. Images were taken at 10× magnification. (B) Illustration of EB-replating assay. (C) SSEA1 staining following EB-replating assay using a 10x lens magnification.

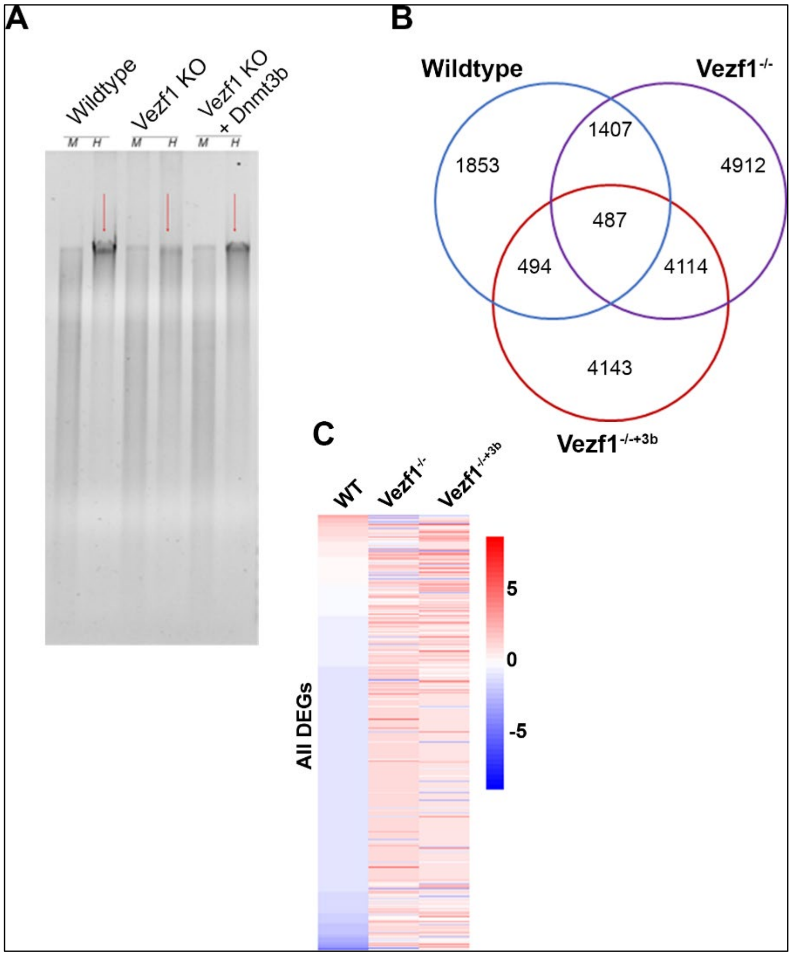


**Figure S2: DNMT3B rescue in *Vezf1*$^{-/-}$ cells**.

(A). Image of an agarose gel showing digested patterns of DNA from WT, *Vezf1*$^{-/-}$, and *Vezf1*$^{-/-+3b}$ treated with MspI (M) and HpaII (H). The ectopic Dnmt3b expression rescued global methylation in *Vezf1*$^{-/-}$ cells, as shown by the similar HpaII digestion patterns for *Vezf1*$^{-/-+3b}$ and WT ESCs (B). Venn diagram showing the percentage of differentially expressed genes (DEGs) in WT, *Vezf1*$^{-/-}$, and *Vezf1*$^{-/-+3b}$ cells post-differentiation (C). Heatmap showing relative gene expression between undifferentiated and D2-differentiated cells. Most genes are downregulated in wild-type cells compared to *Vezf1*$^{-/-}$ mutants.

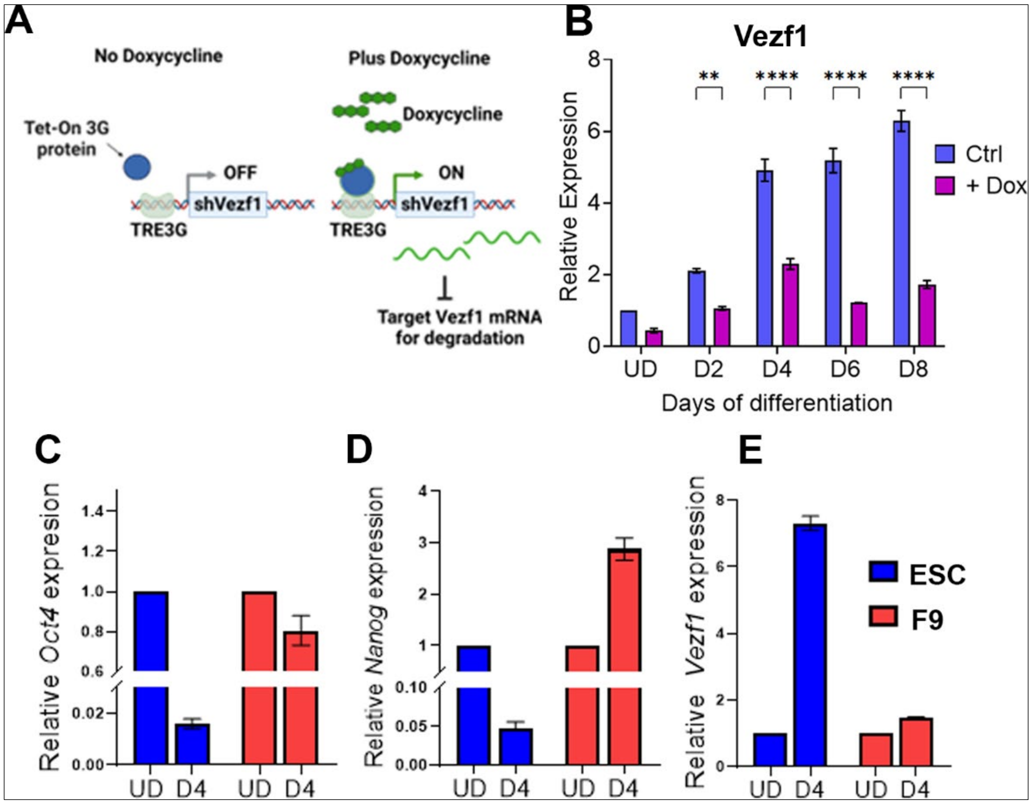


**Figure S3. Modulation of *Vezf1* levels affects PpG expression**

(A) Schematic of Vezf1 knockdown via doxycycline treatment. (B) RT-qPCR showing Vezf1 expression in WT and Vezf1sh- cells on days 3, 5, and 7 post-differentiation, with undifferentiated cells set to 0. (C, D, E) RT-qPCR showing expression of (C) Oct3/4, (D) Nanog, and (E) VEZF1 in undifferentiated ESCs and F9, and on day 4 post-differentiation, with expression in undifferentiated (UD) cells set to 1.

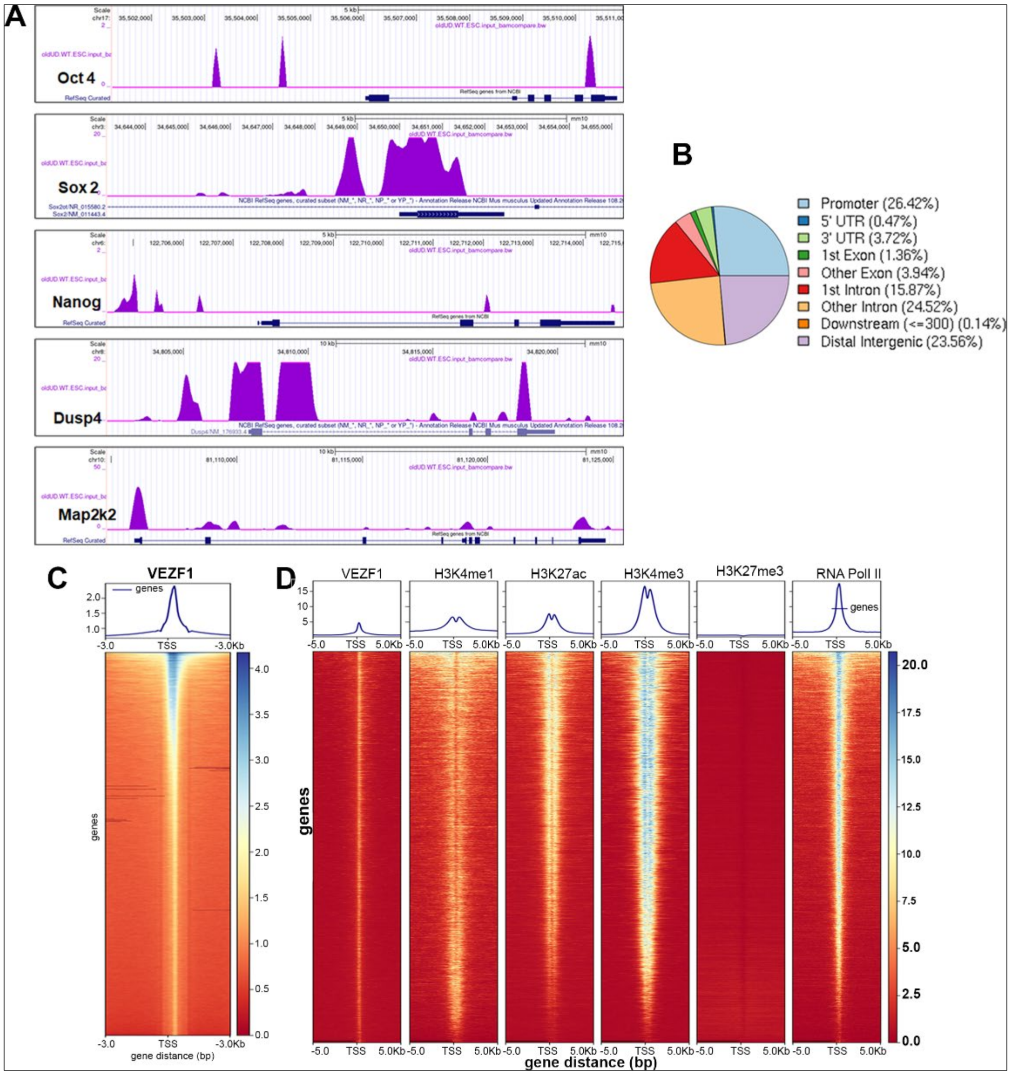


**Figure S4: VEZF1 binding sites are present near active regulatory elements.**

(A) UCSC genome browser tracks of VEZF1 ChIP-seq showing enrichment at pluripotency and MAPK pathway genes, using the mm10 mouse genome as the reference. (B) Distribution of VEZF1 binding sites in the K562 genome. (C) Profile plot and heatmap showing genome-wide VEZF1 binding around the TSS in K562 cells. The scale bar indicates the intensity of Vezf1 enrichment. The profile plot and heatmap were generated using DeepTools. (D) The heatmap shows Vezf1 binding across publicly available ChIP-seq datasets for histone modifications and RNA Pol II in K562 cells.

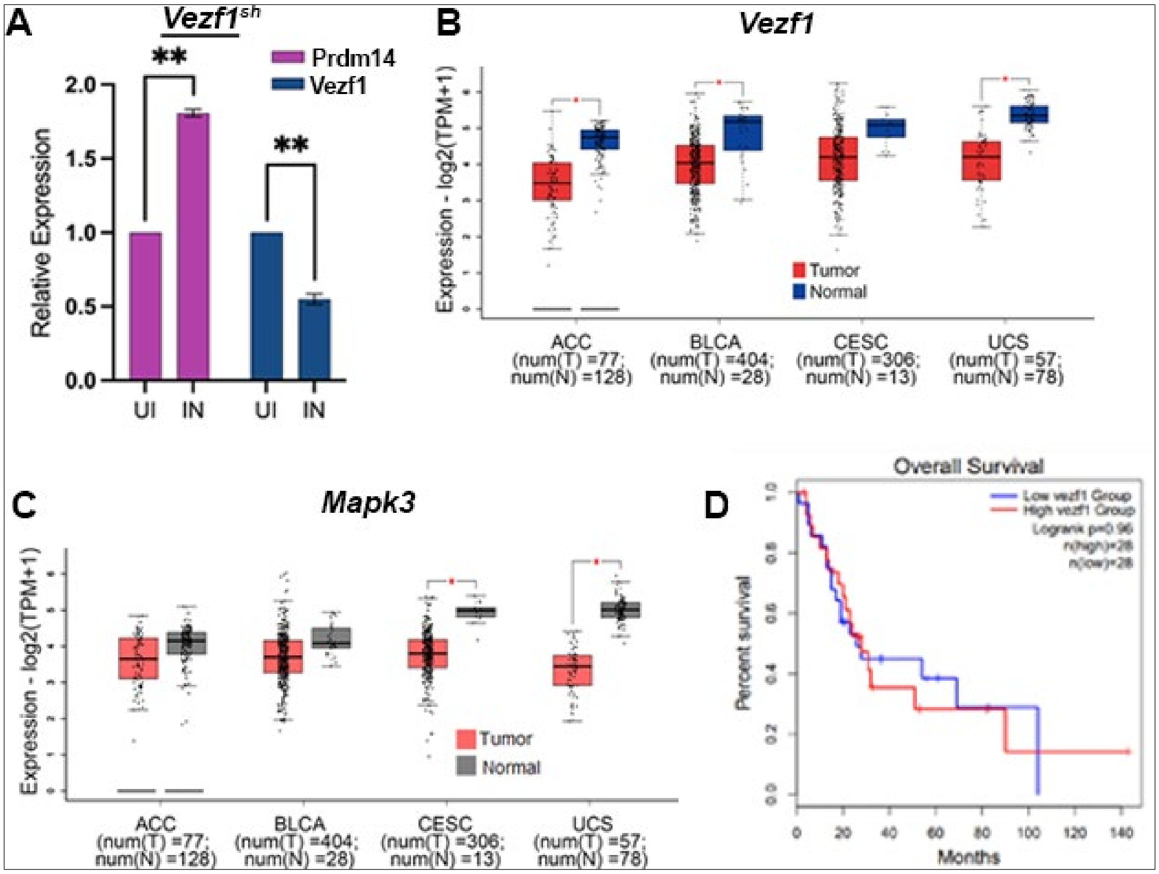


**Figure S5. Differential gene expression analysis in WT and *Vezf1-depleted* cells.**

(A) RT-qPCR showing expression of Prdm14 and *Vezf1* in *Vezf1*$^{sh}$ cells uninduced (UI) or dox-induced (IN). Fold change is normalized to UI set to 1. (B, C) Box plots of RNA-seq data showing the expression of (B) *Vezf1* and (C) *Mapk3* in tumors compared to normal cells from various cancers obtained from the TCGA and GTEx datasets. Box plots were obtained from Gepia2. (D). Survival analysis of UCS patients expressing low *Vezf1* (Blue) or high *Vezf1* (Red).

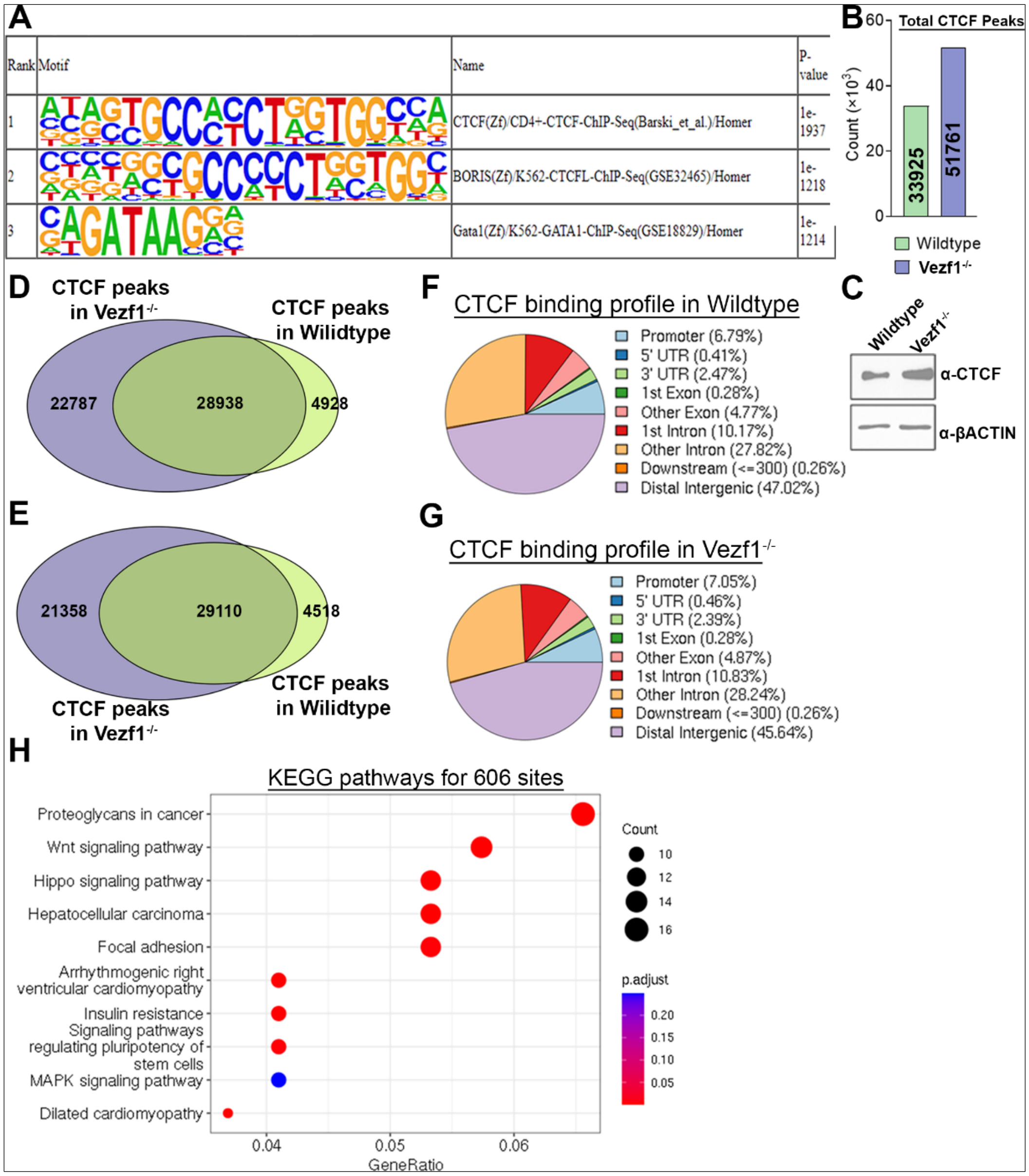


**Figure S6. CTCF binding profile in WT and *Vezf1$^{-/-}$* cells.**

(A) The CTCF motif is enriched in VEZF1 binding analysis following Homer motif analysis. (B). Number of CTCF sites in undifferentiated WT and *Vezf1$^{-/-}$* cells. (C) Western blot showing the expression of CTCF in WT and *Vezf1$^{-/-}$* cells. B-actin was used as a loading control. (D, E) The Venn diagrams show the overlap between VEZF1 and CTCF sites in WT and *Vezf1$^{-/-}$* cells, using a maximum gap of 1000 bp (D) or 2000 bp (E) between binding sites. (F, G) Genomic distribution of CTCF binding sites in WT and *Vezf1$^{-/-}$* cells. (H) KEGG pathway enrichment analysis of 606 CTCF sites in *Vezf1$^{-/-}$* cells that overlap with VEZF1 binding sites.
